# Degradation of sulphonated mono and di-azo dye as the sole carbon source in *Serratia marcescens*: Insights from combined wet and dry lab analysis

**DOI:** 10.1101/2023.06.30.547171

**Authors:** Zarrin Basharat, Azra Yasmin

**Affiliations:** Alpha Genomics (Private) Limited, 45710-Islamabad, Pakistan; Microbiology & Biotechnology Research Lab, Department of Biotechnology, Fatima Jinnah Women University, Rawalpindi 46000, Pakistan

**Keywords:** Azoreductase, *Serratia marcescens*, Methyl orange, Congo red, Docking, Simulation

## Abstract

The high production volume of azo dyes for manufacturing and treating various consumer products leads to deleterious environmental consequences. Bacterial agents present in the environment can degrade these dyes. We, hereby, report the isolation, decolourization and degradation of a mono (Methyl orange) and di-azoic (Congo red) compound of this class of dyes by a versatile bacterium *Serratia marcescens*. Our isolate showed the capability of sulphonated azo dye utilization/degradation i.e. Methyl orange and Congo red usage, with no inhibitory effects on its growth in minimal medium. The calorimetric analysis showed 80.83% decolourization of Methyl orange and 92.7% decolourization of Congo red after 7 days of incubation in a shaking incubator at pH: 7 and temperature: 37 °C. An azoreductase enzyme of ∼25 KDa was detected after SDS-PAGE analysis. Quantitative and qualitative testing of the degradation phenomenon was followed by *in silico* analysis. Structural modeling followed by molecular docking in Molecular Operating Environment revealed numerous residues involved in binding and assisting degradation. Changes in the apo, holo, and dye-bound enzyme energy profiles were also observed. This is the first study reporting the capability of *Serratia marcescens* to use azo dyes/sulphonated azo dyes as the sole carbon source and the detailed computational analysis of the degradation phenomenon. We hope that these findings will be of use to environmental scientists, aid in better dye-degrading mutant creation to help craft future remediation strategies for sulphonated azo dyes.

## 1. Introduction

The lucky discovery of azo dyes by Sir William Henry Perkin in 1853, opened a new avenue for synthetic coloring [1-2]. Based on the presence of single, double, and triple–N=N– bond, azo dyes are termed mono-azo dyes, di-azo dyes, and tri-azo dyes [3]. Besides, some azo colorants own auxochrome functional groups like sulphonic acid, carboxylic acid, hydroxyl groups, and amino groups. These auxochromes do not produce color but only alter solubility, intensity, and the wavelength of the absorbed light [4]. Phenolic compounds chiefly form azoic dye constituents [5]. Azo dyes are extensively used in numerous industries for dying textiles, leather, plastics, etc [6]. These dyes and their residues have become a potential threat to human and ecological health in complex detrimental ways owing to dye recalcitrance to degradation [7]. The estimated release of azoic colorant during the dyeing procedure into the water is approximated to be 10–50% [8]. Extensive usage of these dyes has led to serious pollution problems.

Accumulation and ingestion of azo dyes by aquatic organisms in water bodies leads to deleterious consequences like teratogenicity [9], biomagnification, etc [10]. Mutagenic, cytotoxic, and genotoxic effects of an azo dye investigated by Tsuboy and his colleagues proved that azo dyes kindle DNA fragmentation, micronuclei formation and augment the human hepatoma (HepG2) cell apoptotic index [11]. The toxicity of several of these dyes has led to their ban by the European Union [12], California (The safe drinking water and toxic enforcement act of 1986 i.e. California proposition 65), and other nations [13] but their production and use persists in many parts of the world due to their low manufacturing cost and several other desirable characteristics.

Microbial or enzymatic decolourization of azo dyes is a cost-effective and eco-friendly alternative to chemical treatment method [14]. The distinctive metabolic aptitudes of certain microbes are momentous for bioremediation purposes as these can be utilized for recalcitrant and hazardous environmental pollution cleanup. Oxygen utilizing bacterial strains may utilize azo dyes as a sole carbon source, whereas azoreductase does the job in other [15]. Azoreductase (EC 1.7.1.6) is vital to the degradation of the azoic bond using the ping-pong mechanism and exists in several microorganisms [16]. Here, we report dye decolourization and an azoreductase (AzoR) encoded by *Serratia marcescens*, a bacterium isolated from the effluent of chemical industries that can utilize azo dye as a sole carbon source. This is a gram-negative, facultative anaerobe and known for its ability to degrade various organic compounds. However, to the best of the authors knowledge, analysis of its AzoR enzyme for azo dye (Congo red and Methyl orange) decolourization had not yet been attempted. By focusing on this specific bacterial strain, we contribute to the understanding of its potential in azo dye degradation and expand the knowledge base in this field. We also mined the sequence of this enzyme from the NCBI database and modeled it. This was followed by molecular docking and dynamics simulation analysis for the explanation of the particular biological activity. A combined wet and dry lab experimentation can aid in unearthing the role of catalytic residues and compounds in substrate specificity and activity, which in turn can be used for designing improved mutants and recombinants for enhanced dye degradation. Hence, this work explores the potential of *Serratia marcescens* for azo dye degradation and provides a detailed understanding of the degradation mechanism through enzyme characterization and computational analysis.

## 2. Material and methods

A heavy metal resistant, plant associated *Serratia marcescens* isolated from chemical industrial effluent (Accession no: KJ729142.1), was obtained from the microbial culture collection of our lab. It was initially identified through 16s rRNA sequencing.

### 2.1. Screening of bacteria for dye degradation/utilization capability

Luria-Bertani medium and Bushnell Hass medium were used for primary [17] and secondary screening [18] of dye degradation capability of *Serratia marcescens*. Congo red (CR) (C.I. 22120) and Methyl orange (MO) (C.I. 13025) were purchased from Sigma Aldrich. All reagents used were of analytical grade. The stock solution of MO (0.5 mg/100ml water) and CR (1 mg/90 ml water+10 ml ethanol) was prepared by dissolving the solid chemicals in autoclaved distilled water. The percent decolourization was calculated by the formula [19-20]:

Percent decolourization = (Optical density of control – Optical density of sample) ×100 % / (Optical density of control)

### 2.3. Fourier transform infrared spectroscopy analysis

A 3:1 ratio of Potassium bromide vs pellet of the bacterial degraded product was taken and samples then were ground using mortar and pestle to fine particles and pellets were prepared by establishing a pressure of 100 kg/cm2 (1200 psi). Infrared spectra were obtained by scanning the prepared pellets with a spectrometer (Schimadzu). The spectrum of air was recorded as background and subtracted automatically by using appropriate software. FTIR spectra of degraded dye samples were recorded in the 4000-400 cm^-1^ region [21] at room temperature and the spectrum obtained was overlaid with the dye control and interpreted.

### 2.3. Enzyme extract preparation

The cell-free extract was prepared for activity assay following Aftab *et al*. [22] with necessary modifications. Bacterial cultures were grown in the 100 µg azo dye supplemented with a minimal salt medium for 96 hours at 37 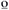C. Cells were harvested by centrifugation for 15 minutes at 9000 rpm. The pellets were then washed twice with 20 mM potassium phosphate buffer (pH:7), frozen at -20 °C, thawed, and then suspended in 10 ml of the same phosphate buffer. Cells were disrupted by the beat-beater method [23]. Universal bottles containing the buffered pellet were supplied with few glass beads and 30 seconds of agitation on vortex was followed by 30-second placement on ice. The process was repeated four times. The homogenate was then centrifuged at 8000 rpm at 4 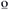C for 20 minutes and the supernatant was used as a crude extract. This was purified by the ammonium sulfate precipitation method followed by dialysis. Solid ammonium sulfate was added to the crude enzyme solution at a concentration of 60%. After dissolving salt in crude enzyme, the solution was placed in cold overnight and then precipitates were collected by centrifugation at 10,000 rpm for 15 minutes. Precipitates were dissolved in the 20 mM phosphate buffer of pH 7 and kept at 4 °C.

A Quantitative Azor assay was then carried out [22] in glass cuvettes with 1ml reaction volume. 400 µL of 20 mM phosphate buffer (pH 7.0) mixed with 200 µL protein sample and 200 µL dye (25 µmol/ml), 200 µL of co-enzyme Nicotinamide adenine dinucleotide hydride (NADH) (1 mM) was monitored spectrophotometrically at 464 nm (for MO) and 498 nm (for CR).

### 2.4. SDS PAGE analysis

SDS page was carried out following the method of Laemmli [24]. 1 ml culture was collected (purified and crude) and cells were prepared for analysis by adding a final 1 X concentration of SDS-PAGE loading buffer, boiling the sample for 15 min, and loading 20 µl onto a 12.5% SDS-PAGE gel [25]. The gel was stained by Coomassie brilliant blue R-250. The broad range protein ladder was used as a marker for the elucidating weight of AzoR.

### 2.5. Structure modeling and dynamics simulation

For computational analysis of the AzoR sequence, the 25 kDa AzoR was obtained from NCBI. This was done by the basic local alignment of 16s rRNA of our strain with fully sequenced genomes of *Serratia marcescens* (encoding AzoR). The one with the least E-value and highest similarity (Accession no: ETX40909.1) was procured for dry lab analysis. The structure was determined [26] by homology modeling using iterative threading assembly simulations in the I-TASSER [27-28]. The structure was then simulated for 10 ns [29] using CABS-Flex server [30]. This was done to visualize the flexibility and stability of the protein chain.

### 2.6. Molecular docking analysis

The substrate molecules Congo red and Methyl orange were downloaded from the PubChem database [31] of NCBI. Open Babel [32] was used for importing SDF files for format conversion and optimization. Profix and TINKER program was run to fix structural defects such as missing atoms of AzoR. The optimized receptor (protonated, energy minimized) and ligand molecules were used for the docking study. Flavin mononucleotide (FMN), NADH, MO, and CR substrate molecules were positioned in the binding pocket by scanning the whole protein surface and flexibly docking the ligands with AzoR in Molecular Operating Environment software. The Triangle Matcher placement method was used for docking. London and affinity dG were used for scoring and rescoring of predicted poses [33]. The top ten structures were retained and the one with the lowest S value was chosen for analysis. Interactions were then visualized. Various energies (angle bend, van der Waals interaction, out-of-plane, bond stretch, dihedral rotation, etc) of ligand-bound and unbound state of AzoR were also studied. The sum of these energies i.e. total potential energy of the system was also recorded.

## 3. Results and Discussion

Genotoxic azo-benzene mediated dye pollution is a threat to both human health and the environment. We are in an urge for fast screening of bacterial varieties with an aptitude for pollutant degradation and hence, find new strains with better degradation capabilities. We explored the potential of *Serratia marcescens*, isolated from chemical industrial effluent, and exploited it for the degradation of sulphonated azo dyes MO and CR.

### 3.1. Growth in MO and CR supplemented media

The plate and quantitative dye reduction assay were carried out for the strain, showing a high rate of decolourization (Fig. 1). The bacterium was tested for dye tolerance up to 250 µg/ml and showed growth at all tested concentrations. Previously, *Serratia marcescens* has been reported to degrade azo dyes (other than sulphonated dyes) but at a lower scale [34]. The degradation activities were attributed to the manganese peroxidase and laccase enzyme. In a study by Mahmood *et al*. [35], another specie of the genus *Serratia* i.e. *Serratia proteamaculans* was reported to completely degrade a sulphonated azo dye (Reactive black 5) in 12 hrs. A molybdenum reducing *Serratia marcescens* could decolorize less than 20% of MO and CR in 48 hrs [36]. Our strain showed way higher decolourization of MO and CR in 48 hrs (Fig. 1).

**Fig. 1.**
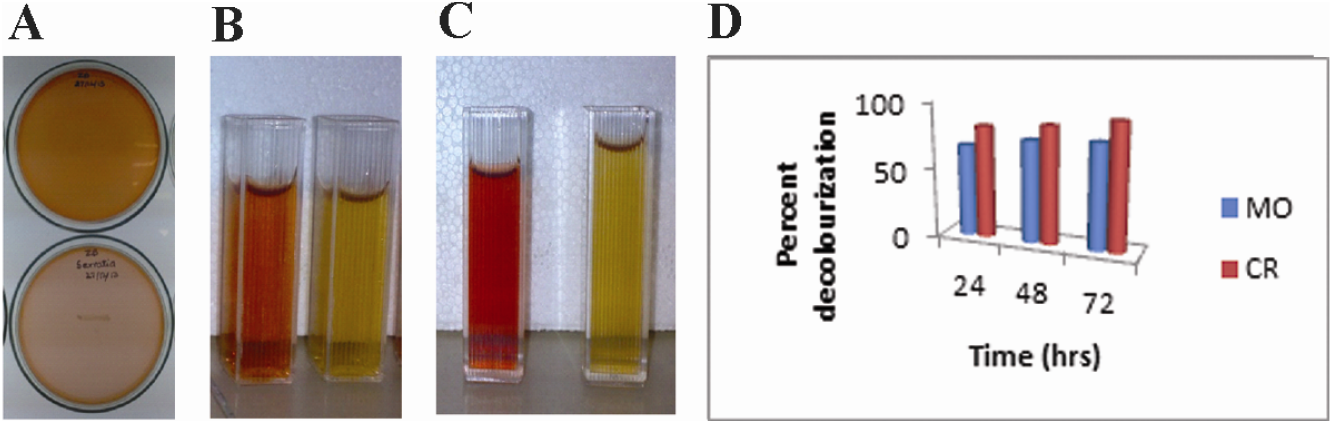
(A) Growth and decolourization of the 50 µg CR dye supplemented BH media after 7 days. The upper Petri plate shows the control. (B) 24 hour decolorized MO, (C) CR in LB medium with controls. (D) Percent decolourization for *Serratia marcescens* in LB dye supplemented media.

### 3.2. Use of MO and CR as the sole carbon source

*Serratia marcescens* could utilize the azo dyes as the sole carbon and energy source. The decolourization percentage was calculated for the strain in liquid BH medium after 96 hours of incubation in the shaking incubator. The decolourization in BH medium for MO after 7 days placement in shaking incubator was 80.83% and for CR 92.7%. Decolourization was more for diazo sulphonated dye as compared to the mono azo dye. It might be due to the structure-activity relation of the dye and the decolourizing enzyme pocket. The growth optical density (O.D.) increased with time and decolourization was dependent on it.

Bacteria capable of aerobic decolourization and mineralization of dyes, especially sulphonated azo dyes, have proven difficult to isolate and they need to be specially adapted for this purpose [37-38]. Formerly, a bacterial strain *Hydrogenophaga palleronii* S1 capable of growing on azo dye as the sole carbon and energy source was obtained by continuous adaptation with dye. The strain could grow cleaved 4-carboxy-4′-sulfoazobenzene reductively under aerobic conditions [39]. Dehydrogenases, menaquinones, and cytochromes were found to be essential electron transfer components for azo reduction during the growth of *Shewanella decolorationis* S12 using the azo compound as the sole electron acceptor [40]. Bheemeraddi *et al*. [41] demonstrated that rapid dye decolourization takes place in the presence of glucose and yeast extract as compared to starch, lactose, sucrose, and other nitrogen sources like peptone, beef extract, potassium nitrate, and sodium nitrate. Organic nitrogen sources might be responsible for NADH regeneration acting as an electron donor in azo bond reduction.

This type of decolourization is rare to find in bioremediation studies but recently, Manogaran *et al*. [42] reported decolourization of a sulphonated bis azo dye, Reactive red 120 by a consortium of bacteria (including *Serratia marcescens*). The consortium required prolonged acclimation for better efficiency or the decolourization was low. Nevertheless, *Serratia marcescens* coupled with *Pseudomonas aeruginosa* and *Enterobacter* sp. could mineralize the dye completely. Several co-substrates or trace elements could enhance decolourization but the addition of yeast showed the opposite effect. It is contemplated that the synergistic effect of the bacterium in consortium might have a beneficial impact on each other to attain the decolourization. Our isolate of the *Serratia marcescens* showed the capability of utilizing azo dyes as a sole source of electron acceptors and with minimal nutritional requirements but it is contemplated that carbon and nitrogen source is imperative to augment the pace of bacterial biodegradation activity. This is the first report of the capability of the axenic culture of *Serratia marcescens* to use azo dyes (sulphonated azo dyes) as the sole carbon source.

### 3.3. FTIR analysis

FTIR analysis of MO control showed specific peaks at 3100 cm^-1^ (asymmetric C-H stretching vibration), 3060 cm^-1^ (=C-H stretching vibration), 2112 cm^-1^ (C-C and CN stretching vibrations), 1700 cm^-1^ (C=C stretching vibration),1624 cm^-1^ (C=C stretching vibration), 1610 cm^-1^ (C=C stretching vibration), 1585 cm^-1^ (C=C stretching vibration), 1500 cm^-1^ (C-H symmetric deformation vibration), 1385 cm^-1^ (symmetric C-H deformation vibration), 1186 cm^-1^ (C-H skeletal vibration), 1050 cm^-1^ (C-H_3_ rocking vibration), 1010 cm^-1^ (C-C skeleton vibration), 990 cm^-1^ (C-C skeleton vibration), 845 cm^-1^ (C-H vibration), 800 cm^-1^ (aromatic C-H out-of-plane deformation vibrations), 720 cm^-1^ (CH^3^ rocking vibration), 655 cm^-1^ (C-H wagging vibration), 642 cm^-1^ (Ring deformation), 595 cm^-1^ (C-C skeletal vibrations, 510 cm^-1^ (C-C skeletal vibration), 495 cm^-1^ (skeletal vibration), 410 cm^-1^ (skeletal vibration), 445 cm^-1^ (skeletal vibration).

*Serratia marcescens* degraded MO product showed peaks at around 2900 cm^-1^ (C-H stretching vibration), 3100 cm^-1^ (N-H stretching vibration-associated to =C), 2400 cm^-1^ (asymmetric N-H stretching vibration),1700 cm^-1^ (C=C stretching vibration), 1600 cm^-1^ (C-H vibration),1506 cm^-1^ (N-H deformation vibrations), 1200 cm^-1^ (C-H skeletal vibration), 1050cm^-1^ (C-H symmetrical deformation vibration), 831 cm^-1^ (C-H out-of-plane deformation vibration), 770 cm^-1^ (N-H out-of-plane bending vibrations).

FTIR analysis of CR control showed specific peaks in the fingerprint region 1650 cm^-1^ (for amide functional group), 1300 cm^-1^ N=N symmetric stretching vibration of azide group, 1310 cm^-1^ (ring structure vibration), around 1400 (napthalene ring), 1505-1550 cm^-1^ (C=C vibration in the naphthalene ring), 1530 cm^-1^ (N=N stretching vibration), 1080 cm^-1^ (C-N stretching vibration of amine), around 990 cm^-1^ (S=O stretching vibration), 1110 cm^-1^ (Ring vibration) including C-S and symmetric SO_3_ stretching vibration, 800 cm^-1^ (C-C skeletal vibration), 660 cm^-1^ (C-S stretching and vibration). For *Serratia marcescens* degraded MO (Fig. 2), peaks became visible at around 2328 cm^-1^ (N-H stretching vibration), 2922 cm^-1^ (C-H stretching vibration), 2900 cm^-1^ (C-H stretching vibration), 2380cm^-1^ (N-H stretching vibration), 1707 cm^-1^(C=C stretch), 1690 cm^-1^ (C=C stretch), 1750 cm^-1^ (C=C stretch), 1590 cm^-1^ (N-H bending vibrations), 1300 cm^-1^ (N=N deformation vibration for azo group), 1090 cm^-1^ (C-H symmetrical deformation vibration), 1200 cm^-1^ (C-H skeletal vibration) and 790 cm^-1^ (C-C skeletal and C-H out-of-plane deformation vibration for aromatic group).

**Fig. 2.**
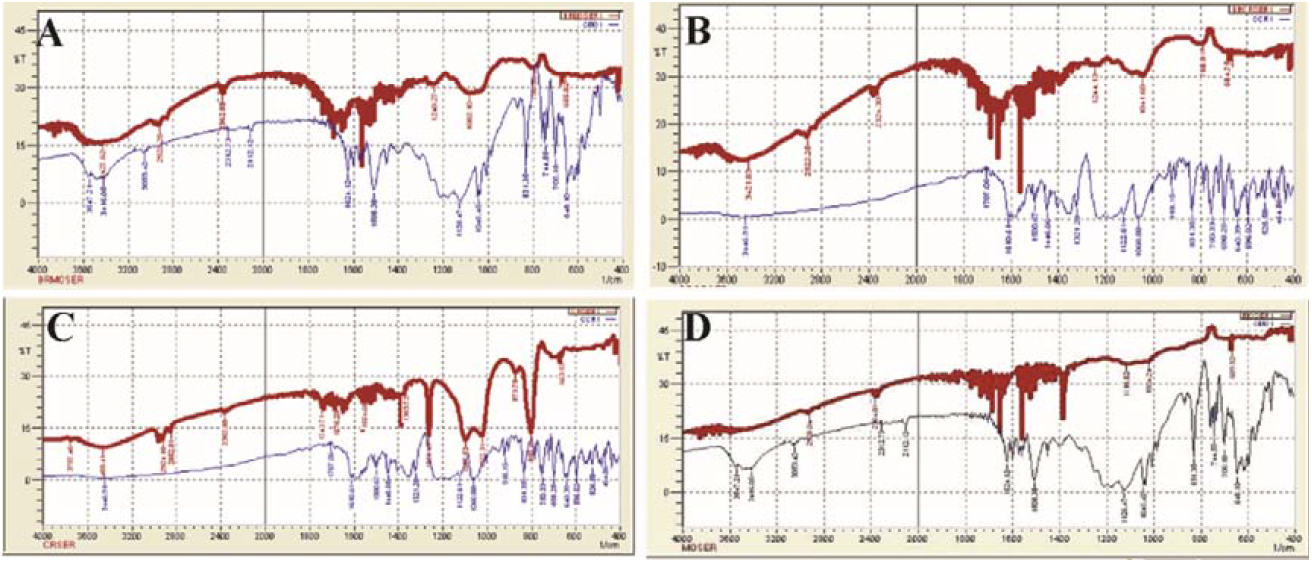
FTIR spectra of degraded dye products for (A) FTIR of degraded MO in LB-medium, (B) FTIR of degraded CR in LB-medium (C) FTIR of degraded MO in BH-medium (D) FTIR of degraded CR in BH-medium.

In IR spectra of MO and CR degradation reported by Quan *et al*. [43], non-uniform vibrations of 1584 cm^-1^ (for -N=N-of azo dye), 1450 and 1510 cm^-1^ (benzene skeleton vibration), 1180 cm^-1^ (benzene ring in-plane bending vibration), 698 and 536 cm^-1^ (outward flexural vibration of benzene ring) were detected. Degradation of the dyes caused a lot of peaks to either weaken or disappear, indicating the occurrence of a decolourization reaction. Krithika *et al*. [44] report that the FTIR spectra of controls showed several peaks in the 3200-3500 cm^-1^ region (N–H and O–H stretch). After the breakdown of dyes, absorption decreased for this region. New bands were formed in the carbonyl region (1401.07–1072.83 cm−1), attributed to amine groups. Our strain also showed amine formation (N-H deformation, bending, or stretching vibrations) in both spectra recorded for degraded MO and CR. 1506 cm^-1^.

### 3.4. AzoR characterization

Enzymatic bioremediation is a potentially rapid method involved in the metabolic and co-metabolic transformation, detoxification, or degradation of toxic dye [45]. The preliminary strategy of the azo dye biodegradation is decolourization, where - N=N-is cleaved in a reductive manner. This step is catalyzed by the enzyme AzoR [46]. A 25 KDa AzoR enzyme band was observed in our isolate of *Serratia marcescens*, after SDS PAGE analysis. Eslami *et al*. [47] previously reported a 22 Kda AzoR from a halophilic bacterium whereas a 29 KDa AzoR was isolated by Nisar *et al*. [48] from a *Staphylococcus* sp. Dong *et al*. [49] reported many copies of NAD(P)H-dependent AzoR in *Serratia* sp. S2. They were found to have a role in chromium remediation. Three copies of AzoR enzyme were found to be encoded by a *Serratia* sp. when the whole genome was sequenced by Basharat et al. [50].

This proves that *Serratia marcescens* encodes AzoR enzyme that can be used for remediation in case of excessive dye quantity hindering the activity of target molecule or in case of additive induced growth inhibition. Another characteristic of AzoRs is that they produce aromatic amines via reductive cleavage of -N=N-. Our strain also showed a tendency towards amine formation, which was seen in FTIR analysis, so this further strengthens the observation that the decolourization was via AzoR. The feasibility of enzymatic treatment has been demonstrated at laboratory scale in prior studies and we propose that AzoR may be obtained in large amounts from *Serratia marcescens* grown in favorable conditions.

### 3.5. Impact of reducing factors on degradation

Azo dye degradation can be achieved by AzoR because it cleaves the azo bond [46] by specific oxidation or reduction at the expense of a reducing agent. It ensnares azo dyes and acts as an effective dye degrading tool by using the ping pong mechanism; i-e ambushing or locking in the toxic azo dyes in the enzyme, with AzoR being the main entity and using supporting factors (reducing agents) like NAD/NADH/FAD or FADH. Dyes are reduced via a ping-pong bi-bi process which entails two cycles of NAD(P)H-based FMN to FMNH_2_ reduction [51]). During the first phase, the azoic substrate reduces to hydrazine whereas, in the second round, hydrazine is reduced to two amines [52]. AzoR based dye-reduction phenomenon can be divided into two types based on the processing pathway of the enzyme i-e membrane-bound and cytoplasmic AzoR based dye-reduction. Cytoplasmic AzoR enzymes can reduce non-sulphonated azo dyes as they can diffuse through the bacterial cell membrane. For membrane-bound AzoR decolourization, they can use a redox mediator to shuttle the electrons across the membrane barrier in a non-aerobic environment [15]. Azoic dyes comprising sulphonate groups and higher molecular weights are not likely to be transferred across the cell membranes. As a result, the reducing action of the dye is independent of the intracellular dye uptake [53-54]. Reduction in the extracellular environment is based on electron transport and is achieved by the establishment of a link between intracellular electron transport systems of a bacterial cell and azobenzene dye molecules of high molecular weight [15]. It was suggested by Russ *et al*. [55] that bacterial membranes are nearly impermeable to co-factors having flavin, hence, restricting the reduction of sulphonated azo dyes due to equivalent transfer by cytoplasmic flavins. An alternate mechanism for sulphonated azoic-dye reduction in cells with intact membranes occurs due to the genesis of reduced flavins by the cytoplasmic flavin-dependent AzoR [54]. The outer membranous electron transfer apparatus of the bacterial cell wall either makes contact directly with the azo dye substrate or interacts circuitously to the cell surface redox mediator. Redox mediator compounds with small molecular weight can act as electron carriers among the outer membranous NADH-dependent AzoR and azo dye. Bacteria may synthesize redox mediators during substrate processing or these are added superficially [56]. Various researches have proved that the addition of redox mediators accelerates the dye degradation process [56-58]. Our quantitative AzoR assay showed an external addition of NADH/NADH was not necessary for dye degradation, which means it was encoded by the bacterium itself.

### 3.6. 3D structure analysis

Implementation of successful enzymatic bioremediation approaches relies profoundly on intrinsic molecular analysis of the microbial enzymatic structure. Protein structural modeling and docking is now a mature technique and a powerful tool for the understanding and description of protein forms and functions, although careful consideration must be utilized when choosing a template. The protein sequence of *Serratia marcescens* was analyzed computationally to obtain insights about substrate specificity, reaction mechanism, electron transfer, dye positioning, and interaction with amino acid residues of AzoR. Our structure showed more fluctuation at the loop regions as compared to β-sheets and α-helices (Fig. 3). FMN prosthetic group was most probably bound to the enzyme or AzoR was flavin independent, as there was no requirement for the exogenous addition of flavins. This was in correspondence with the performed AzoR assay where no flavin moiety was added externally but decolourization was observed. The motif GXGXXG was absent from our specie. Although *Serratia marcescens* in this study lacked GXGXXG motif but a general β-α-β-α-β structure was present where FMN might bind (Fig. 3). In *Bacillus* sp. AzoR in monomeric form, FMN has been found to be bound in the loop pocket [15, 59]. These insights also lead to the idea that AzoR in *Serratia* sp. might be flavin free. Due to the lack of experimental structures of AzoR specie till todate, we are unable to say conclusively that this might be the case and more studies in this area could help reach valid reasoning. Nevertheless, we did interaction analysis with FMN *in silico* and found several residues to bind with the moiety (section 3.7).

**Fig. 3.**
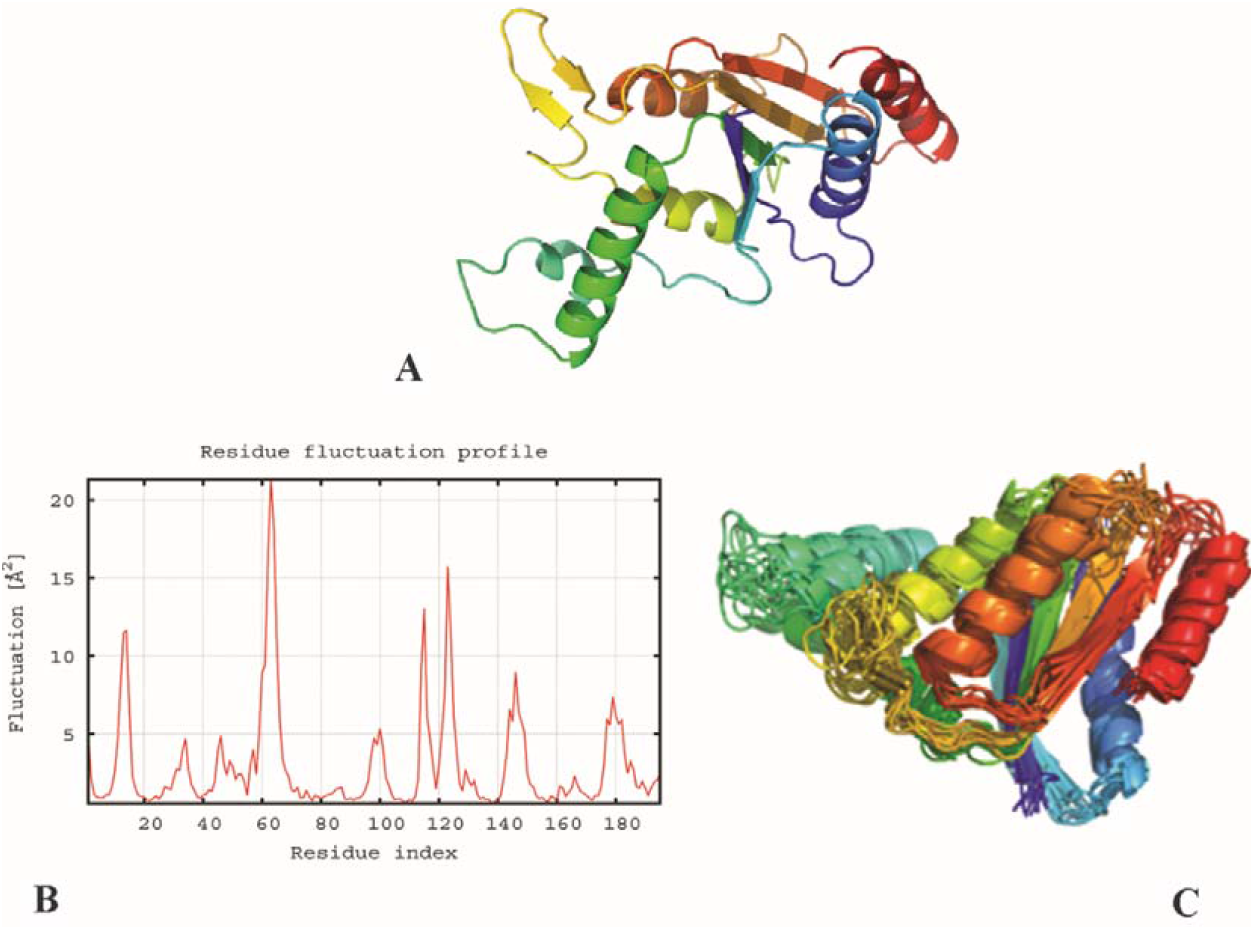
(A) 3D modeled structure of *Serratia marcescens* AzoR. (B) Structural flexibility profile of simulated AzoR, with fluctuations for individual protein residues shown in the red line. The output is based on the all-atom model via trajectory clustering. (C) Refinement of the model and superpositioning (shown in 3D) is centred on the maximum likelihood superposition method of THESEUS (Theobald and Wuttke, 2006).

As far as Rossman binding fold is concerned, Sarkar *et al*. [15] has previously reported five beta sheets and a high percentage of alpha helices in AzoR from *Bacillus* sp. similar to AzoR of our *Serratia* specie. This explains that Rossman binding fold (six parallel beta strands linked to two pairs of α-helices in the topological order β-α-β-α-β) is not universal.

### 3.7. Hydrogen and hydrophobic interactions in dye degradation

A protein is capable of taking complex shapes and its interactions with other proteins or ligands allow it to form versatile conformations. Modeling and abstraction have played a noteworthy role in perceiving protein-ligand associations using computational modus operandi. Molecular docking of the enzyme with various substrates revealed residue and type of interactions present (Fig. 4).

**Fig. 4.**
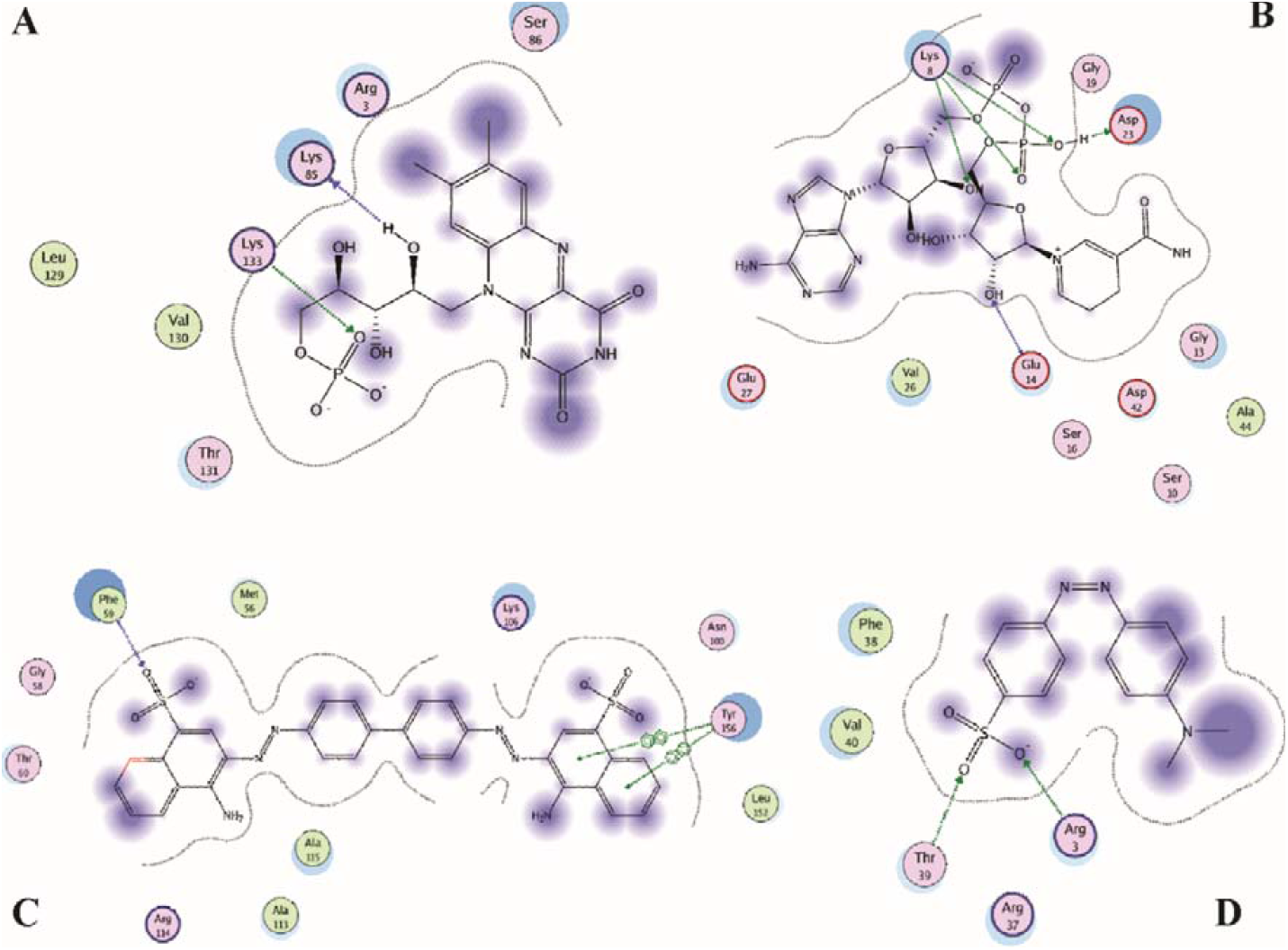
2D representation of (A) FMN-AzoR complex (B) NADH-AzoR complex (C) CR-AzoR complex (D) MO-AzoR complex. 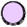 depicts polar residue, 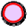acidic,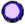 basic, 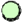hydrophobic, 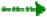side chain acceptor, 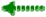side chain donor, 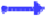backbone donor, 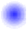ligand exposure, 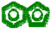arene-arene and black dotted line as proximity contour.

A hydride is typically transferred from NADH to FMN to accomplish the first stage of the reduction reaction. Three hydrogen bonds (two with Lys8 and one with Glu14) and eight hydrophobic interactions (with residue Gly13, Ser16, Gly19, Asp23, Val26, Glu27, Asp42, Ala44) were observed between NADH and AzoR. Cao *et al*. [60] reported slightly different interactions for NADH-AzoR binding in *Shewanella onediensis* MR-1. Ser16 and Ala residue formed hydrogen bonds instead of hydrophobic interactions, as observed in our study. Apart from this study, we could not find reduced NAD interacting residues with azoreductase reported in literature till todate.

For FMN-AzoR interaction, a hydrogen bond was observed between the nitrogen of Lys133 of AzoR and the oxygen atom of FMN. Lys85, Val130, Thr131, and Arg33 form hydrophobic interaction with oxygen atoms of FMN while Ser86 formed a hydrophobic interaction with the carbon atom of FMN. The pattern of FMN binding was compared to Bacillus valezensis [59], and common interacting residues were Arg and Thr (although hydrophobic interactions instead of hydrogen bonding were observed). Hydrogen bond interactions of the *Chromobacterium violaceum* [61], *Bacillus* sp. [62], and *Klebsiella* sp. (PDB ID: 6DXP) were different and entailed only one residue (i.e. Arg) common to that of FMN-AzoR interacting residues of our bacterium.

MO formed two hydrogen bonds (with residue Thr39 and Arg3) and three hydrophobic interactions (with residue Arg37, Phe38, Val40). Six of these residues (Arg, Thr, Phe, Met, Gly, Asn) were common to interactions of AzoR-MO in *Oedogonium subpalgiostomum* AP1 [63]. None of our MO binding residues were similar to that of *Pseudomonas putida* [64]. CR formed one hydrogen bond and ten hydrophobic interactions (with residue Met 56, Gly58, Phe59, Thr60, Asn100, Lys106, Ala113, Arg114, Ala115, Leu152, Tyr156). Two of these residues were common when compared to CR-AzoR interaction of Pseudomonas putida [64].

As azo dyes require polar electron-donating ring substituents (e.g. -OH, -NH_2_, - NHCH_3_, or -N(CH_3_)_2_) so the reactivity of azo dye substrates is determined by their electron densities and redox potentials [65-66]. Punj and John [67] demonstrated that H^+^ bond formation can reduce the electron density around the residue making it more amendable to reductive cleavage. This might explain more degradation of MO as compared to CR by *Serratia marcescens* AzoR because two hydrogen bonds were observed in MO-AzoR interaction while only one hydrogen bond was observed in the case of CR-AzoR interaction. The energy profile of the ligand-bound and unbound states of AzoR also varied (Table 1).

**Table 1.**
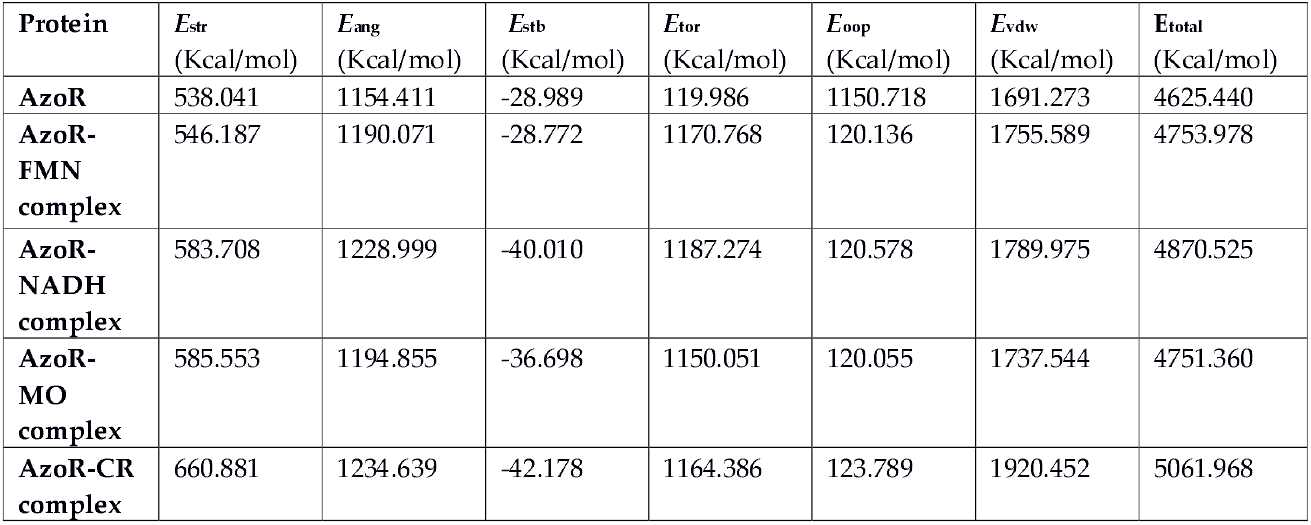
Energy profile of AzoR and its ligand-bound complexes. A potential energy model, equivalently, a forcefield (Amber99 in this case), assigns a potential energy value to a molecular configuration. E_str_ = bond stretch energies, E_ang_ = angle bend energies, E_stb_ = stretch-bend cross term energies, E_tor_ = dihedral rotation energies, E_oop_ = out-of-plane energies, E_vdw_ = van der Waals interaction energies. The potential energy is the sum of interaction energies, depicted as E_total_ in the table.

These results further advance our knowledge on the molecular aspects of the interaction of sulphonated azo dyes to AzoR enzyme in *Serratia marcescens*, depicting the integral role of reductases in environmental cleanup. Thus, this work can also serve as a baseline for the mechanistic understanding of factors affecting enzyme reactivity towards a class of substrates rather than just one single substrate. Future work can be targeted towards the assessment of quantitative structure-activity relationships, incorporating maximum reaction rate and substrate specificity study for bioremediation-need driven practical designing of waste-treatment catalysts.

## 4. Conclusion

This study aimed to present a lucid picture of the relation and communication between FMN, NADH, dyes, and substrate molecule. We were able to identify residues crucial for interaction between AzoR and substrates. Comparison revealed that these were not totally conserved in comparison to the available studies. However, if they are conserved within the genus *Serratia* remains to be elucidated. The acquired information can aid in designing AzoR enzyme mutants for better accommodation of dye pollutants and thus, be of benefit to the community working on enzymatic bioremediation. Additional proposed work is an optimization of AzoR mediated dye degradation studies using response surface models, site-directed mutagenesis for better dye affinity to the enzyme, and reconstruction of azo-dye degradation/metabolic pathways in these organisms. This might be helpful to curb free radical, hydrophobic character, electrophilic species, and arylamine derivative formation leading to interaction with electron-rich sites in DNA, curtailing mutation, and DNA adduct formation responsible for cancer. Also, the development of fluorophores/biosensors for the detection of AzoR activity in bacterial cultures as well as in environmental samples for monitoring and analysis is proposed.

## Declarations

### Funding

This research did not receive any specific grant from funding agencies in the public, commercial, or not-for-profit sectors.

### Conflict of Interest

The authors have nothing to disclose in terms of conflict of interest.

### Ethics approval

Not applicable.

### Consent to publish

All authors consent to be responsible for the published material and have read and approved the content.

### Availability of data and material

This work was done using public data from NCBI. All data and related material is present in the manuscript and accession numbers mentioned at appropriate places. No new data was generated that requires submission in a repository and no plant or animal material was used that requires ethical approval.

### Author contributions

A.Y. conceived and designed the project, provided resources, supervised and edited the paper. Z.B. designed the project, performed the experiments and wrote the paper.

## Notes

### Competing Interest Statement

The authors have declared no competing interest.

